# ChronoClone links temporal clonal behavior to lineage output and mutation responses

**DOI:** 10.64898/2026.11.24.768931

**Authors:** Hiromitsu Harimoto, Weizhong Zheng, Berthold Göttgens, Melania Barile, Nicola K. Wilson

## Abstract

Distinct stem cell clones can yield similar lineage outputs via different developmental trajectories or dynamics and respond differently to perturbation. Here, we introduce ChronoClone, an unsupervised method that groups single-cell clones by time-resolved cell-state composition into metaclones. Benchmarked against simulated ground-truth scenarios, ChronoClone demonstrates high noise tolerance compared to existing methods. Applied to an adult mouse hematopoietic lineage-tracing dataset, ChronoClone showed that clones producing the same lineage mature at different speeds, a difference the original study did not examine. Furthermore, using WT metaclones as a baseline reference for Dnmt3a- and Npm1c-mutant sister-clones, ChronoClone uncovered hidden, clone-specific mutation responses including slowed maturation and lineage composition shifts. By connecting temporal dynamics, perturbation responses, and ancestral molecular states, ChronoClone adds temporal cell-state dynamics into clonal analysis.

**Highlights:**

- ChronoClone classifies stem cell clones by their time-resolved cell-state composition
- Baseline metaclones reveal hidden, clone-specific perturbation dynamics
- Clonal dynamics guide retrospective comparisons of early progenitors’ gene programs
- ChronoClone operates independently of gene expression data, enabling broad application across diverse single-cell technologies and tissues.

**In brief:** Harimoto et al. introduce ChronoClone to organize clonal output by temporal behavior. ChronoClone groups clones with similar temporal and cell-state behaviors into metaclones. The metaclones serve as a reference for examining perturbation responses and enable retrospective comparisons of early progenitor molecular programs. Additionally, ChronoClone is applicable to any longitudinal lineage-tracing data.

## Introduction

Clones producing the same lineage can differ in when their descendants leave early states and which differentiation routes they follow^1–4^. Heritable barcodes combined with single-cell RNA sequencing enable transcriptional state to be connected to clonal fate^5,6^. However, sequencing is destructive, meaning temporal clonal behavior must be inferred from the time resolved composition of sampled descendants. Current computational approaches address complementary aspects of lineage tracing; Cassiopeia reconstructs phylogenies^7^, LineageOT and CoSpar infer ancestor–descendant relationships or transition maps^8,9^, CLADES groups clones and models their differential dynamics^10^, and clone2vec constructs continuous clonal embeddings from cellular expression neighborhoods^11^. Despite these advances, there remains a need for methodologies that group clones directly by how their descendants cell states change over time and can use these groups as a reference in perturbation experiments.

To address this gap, we developed ChronoClone, an unsupervised method that groups clones with similar time-resolved behavior into metaclones. We demonstrate ChronoClone’s utility in three broadly applicable core applications: i) separating dynamic trajectory kinetics from static endpoint outputs, ii) using baseline reference transfers to uncover subtle mutant responses and iii) tracing future dynamic behaviors back to molecular programs in early progenitors.

## Results

### ChronoClone defines metaclones from time-resolved clonal state composition

The input for ChronoClone is longitudinal lineage-traced single-cell data, sample time points, starting population definitions and optional condition labels (Figure 1A). After excluding clones containing cells other than the starting (e.g. earliest progenitor) cell type(s) at the initial time point (Figure 1A i), counts are normalized to clonal cell composition per time point, forming a “clonal” count matrix for downstream analysis (Figure 1A ii).

**Figure 1.**
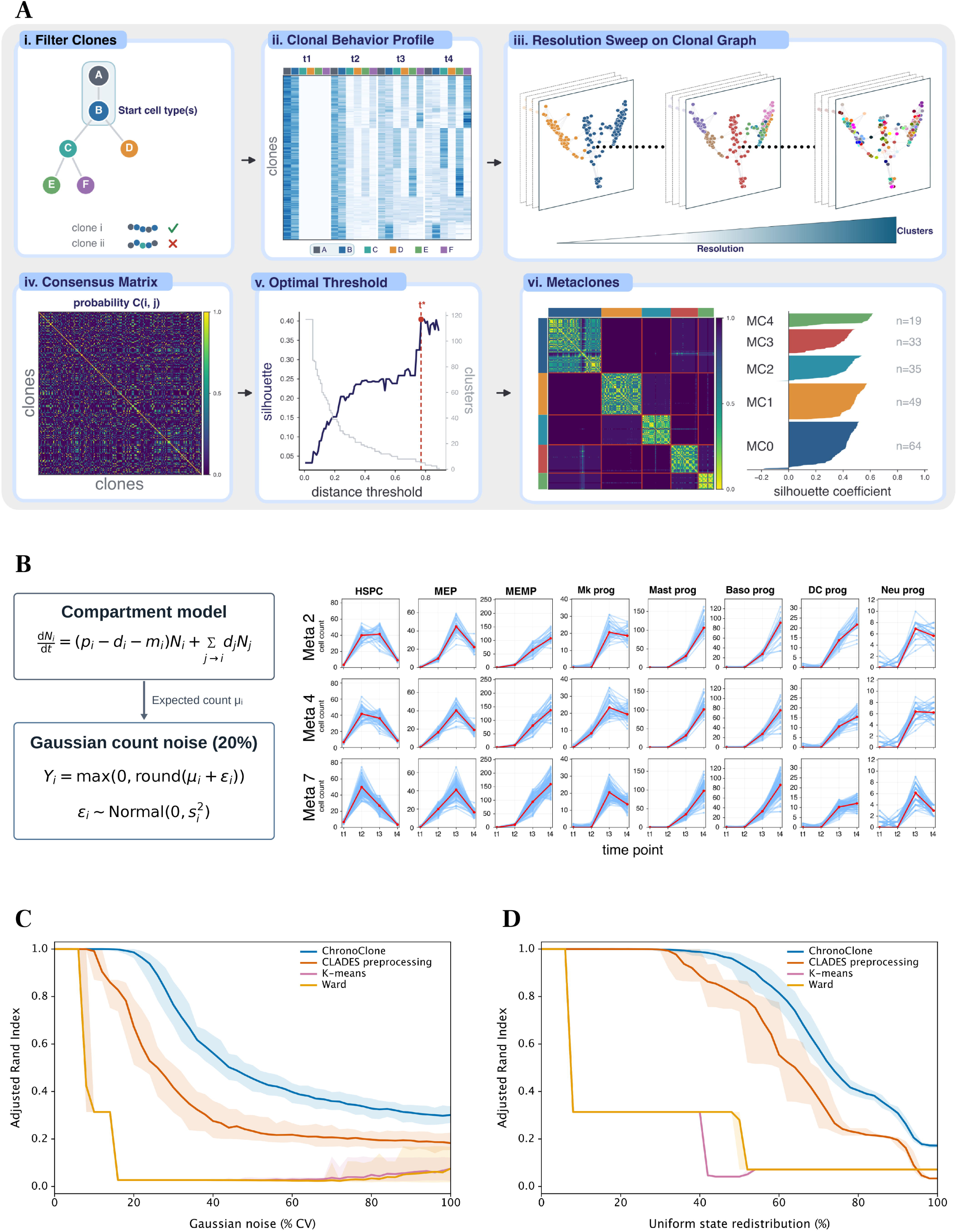
ChronoClone defines metaclones from time-resolved clonal state composition. **A.** ChronoClone workflow: selection of clones that start in the defined starting cell types, cell-state proportions at each time point, Leiden clustering over a range of resolutions, consensus clustering, and choice of the partition with the highest silhouette score. Values are illustrative. Metaclones defined in the reference condition are then kept fixed. **B.** Simulated data. A compartment model with proliferation, differentiation and loss rates defines 12 time-resolved programs over 13 cell states and four time points. Red curves show expected cell counts (μ) for three programs and eight cell states. In the model, N is the abundance of state i, and p, d and m are its proliferation, differentiation and loss rates. Blue curves show counts after adding Gaussian noise at each of the four time points, rounding to the nearest integer and clipping negative values to zero. The noise standard deviation (s) is 20% of the expected count, or one cell when the expected count is zero. In the recovery benchmarks in C and D, noise is applied only at t2–t4. HSPC – hematopoietic stem and progenitor cell, MEP - Megakaryocyte–erythroid progenitor, MEMP - Megakaryocyte–Erythroid–Mast Cell Progenitor, Mk prog – megakaryocyte progenitor, Mast prog – mast cell progenitor, Baso prog – basophil progenitor, DC prog – dendritic cell progenitor, Neu prog – neutrophil progenitor. **C.** Adjusted Rand index (ARI) between the groups found by each method and the simulated programs, as Gaussian noise increases. Lines show the mean and bands the 2.5th–97.5th percentiles of 100 repeats. **D.** As in C, for random redistribution of cells between states (the number of cells per clone and time point is unchanged). All methods are given the same simulated clones and choose the number of groups without access to the true programs.

A neighborhood graph is partitioned using Leiden clustering^12^ (Figure 1A iii) and clonal pair-wise co-clustering frequencies are calculated to summarize agreement among clusters^13^. The resulting consensus probability matrix is then separated into candidate metaclones via hierarchical clustering (Figure 1A iv). ChronoClone retains the partition with the highest silhouette score^14^, therefore, eliminating user bias in the number of metaclones selected (Figure 1A v).

To benchmark ChronoClone against ground-truth, we simulated 517 clones across 12 dynamic time-resolved behaviors using an ordinary differential equation (ODE) model perturbed with Gaussian count variation and random state redistribution, thereby mimicking real-life biological and technical variances such as incomplete- or mis-sampling and imperfect reads (Figure 1B, STAR Methods; Table S1).

ChronoClone maintained a mean Adjusted Rand Index (ARI) of 0.986 under 20% Gaussian noise (Figure 1C) and 0.942 under 50% state redistribution (Figure 1D), outperforming CLADES preprocessing^10^ (ARI 0.673 / 0.800), K-means (0.027 / 0.042) and Ward Clustering (0.027 / 0.280). ChronoClone accurately recovered the 12 ground-truth differentiation programs across moderate noise regimes (Figures S1A–S1F and STAR Methods).

### ChronoClone resolves asynchronous dynamics and lineage choice within similar lineage outputs

We applied ChronoClone to a LARRY (Lineage and RNA RecoverY) lineage tracing dataset of in vitro differentiation of normal hematopoietic stem/progenitor cells^5^ (Figure 2A). ChronoClone processed 569 clones (7,733 cells) spanning neutrophil, monocyte, erythroid/megakaryocyte, basophil and multilineage output clones (Figures 2B, 2D, S2A and S2B; Table S2) and grouped the temporal behaviors into 20 metaclones. Five metaclones (MC1, MC9, MC14, MC16 and MC22) exhibited >93% neutrophil output, yet ChronoClone revealed temporal heterogeneity (Figures 2D and 2E): Day-6 median pseudotime ranged from 0.432 (MC1) to 0.981 (MC14), whilst MC9 failed to reach late granulocytic states despite similar early dynamics to that of MC14 (Figure 2F). Similarly, among monocyte-biased metaclones (>90% output, MC6, MC7, MC8 and MC12) (Figure 2D and 2G), MC7/MC12 exhibited rapid Day 4 maturation (pseudotime 0.478 and 0.460 respectively), compared to MC6/MC8 (0.236 and 0.211 respectively) (Figure 2H and Table S2). Fast- and slow-maturing clones also differed in how their size changed over time (Figure S2C; STAR Methods). They had the same number of sequenced cells per clone at Day 2 (median 1), whereas fast-maturing clones were slightly larger at Day 4 (4 versus 3 cells; P = 0.005), and by Day 6 were much smaller (1 versus 7 cells; P < 0.001 – the majority of MC7 clones were no longer detected). Fast maturation was therefore followed by a decline in clone size, whereas slow-maturing clones continued to grow. Highlighting the ability of ChronoClone to depict distinct behaviors between metaclones, not limited to lineage output but also dynamic temporal behaviors and cell production rates, emphasizing the large amount of biologically meaningful information that can be extracted by jointly considering lineage choice and time- resolved differentiation dynamics.

**Figure 2:**
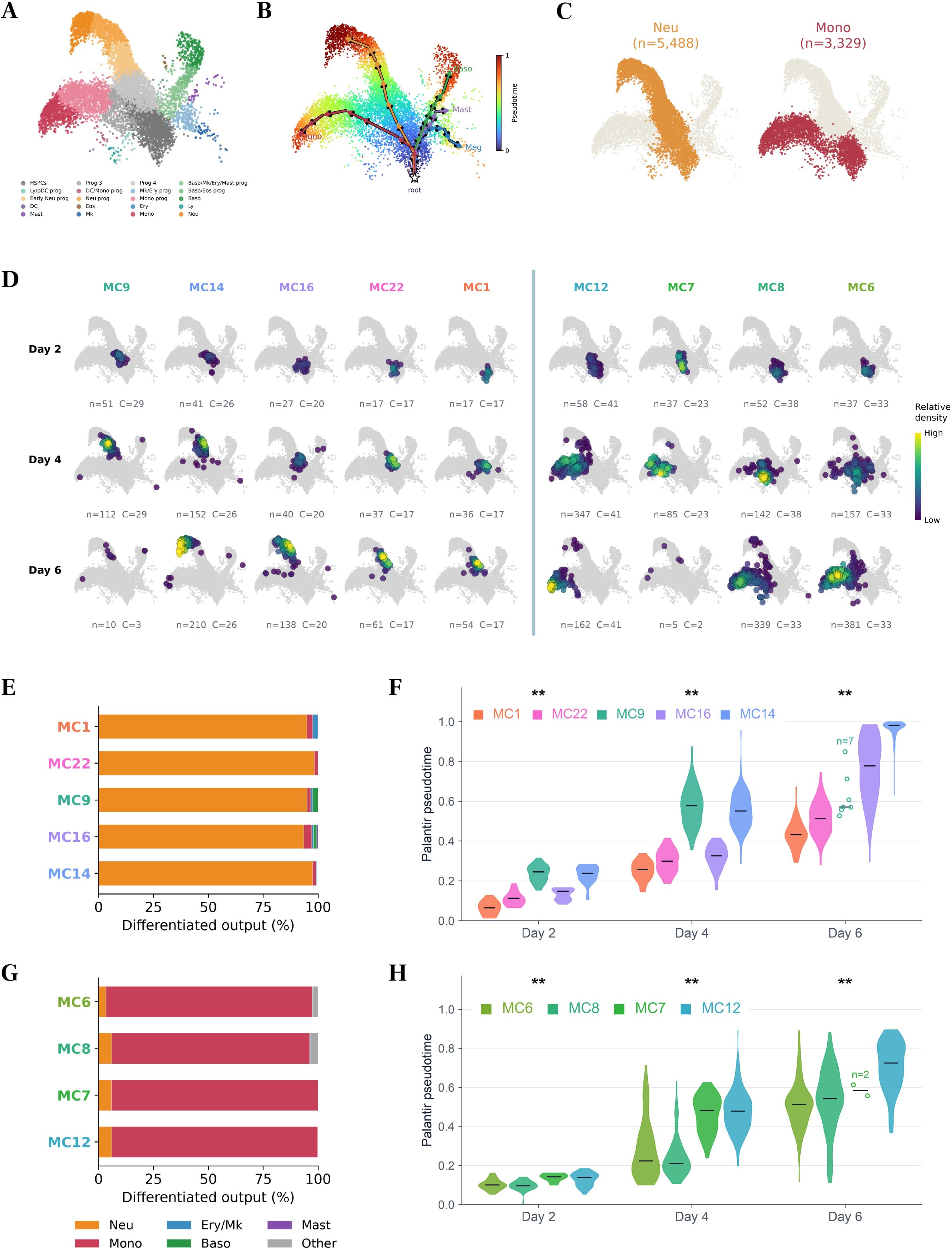
ChronoClone uncovers dynamic differences within metaclones with similar lineage outputs. **A.** LARRY (Lineage and RNA RecoverY) lineage tracing dataset of in vitro differentiation of normal hematopoietic stem/progenitor cells^5^. Cell-state landscape and annotations taken from Gao et al.^10^. HSPC – hematopoietic stem and progenitor cell, Ly/DC prog – lymphoid/dendritic cell progenitor, early Neu prog – early neutrophil progenitor, DC – dendritic cell progenitor, DC/Mono prog – dendritic cell/monocyte progenitor, Neu prog – neutrophil progenitor, Eos – eosinophil progenitor, Mk – megakaryocyte progenitor, Mk/Ery prog – megakaryocyte/erythroid progenitor, Mono prog – monocyte progenitor, ery – erythroid progenitor, mono – monocyte, Baso/Mk/Ery/Mast prog – basophil/megakaryocyte/erythroid/mast cell progenitor, Baso/Eos – Basophil/Eosinophil progenitor, Baso – basophil, Ly – lymphoid, Neu – neutrophil. **B.** Global Palantir pseudotime with segmented trajectories and the root cell marked by a white star. The identified trajectories are shown in C and Figure S2B. Trajectories overlap because a cell can belong to more than one branch. **C.** Identified Neutrophil and Monocyte trajectories, shown on the transcriptomic landscape in A; n denotes number of cells. **D.** Day-2, Day-4 and Day-6 distributions of every neutrophil-output metaclone (left) and every monocyte-output metaclone (right) on the HSPC transcriptomic landscape. Color shows relative density within each metaclone and day; n denotes number of cells and C the number of clones. The remaining metaclones are in Figure S2A. **E.** Differentiated cells of the neutrophil-output metaclones, pooled over the three days (undifferentiated cells excluded). **F.** Palantir pseudotime of neutrophil-trajectory cells in the neutrophil-output metaclones on each day; black bars mark the median. ** q < 0.05, Kruskal–Wallis test of whether the metaclones differ in clone median pseudotime on that day (Benjamini–Hochberg correction over 12 tests; STAR Methods). MC9 has only two clones on the trajectory at Day 6 and was left out of that day’s test. Groups with fewer than eight cells are shown as individual points. **G.** Differentiated output for MC6, MC8, MC7, and MC12 (monocyte producing metaclones) shown as in E. **H.** Palantir pseudotime of monocyte-trajectory cells in the monocyte-output metaclones, shown as in F. ** q < 0.05, Kruskal–Wallis test of whether the metaclones differ in how far their clones have progressed along the trajectory on that day. MC7 has only two clones on the trajectory at Day 6 and was left out of that day’s test.

For multilineage metaclones (MC0, MC2, MC3, MC4, MC5, MC11 and MC18), ChronoClone uncovered time-dependent lineage output (Figure 3A). MC0 gave rise to myeloid cells that differentiated and matured during in vitro culture. MC5 on the other hand, whilst producing cells of the myeloid lineage favored production of neutrophils, whereas MC11 and MC18 retained mostly early-state progenitors together with multilineage output. MC2 mainly produced basophils and neutrophil progenitors, whereas MC4 maintained a multilineage output at both Day 4 and Day 6.

**Figure 3:**
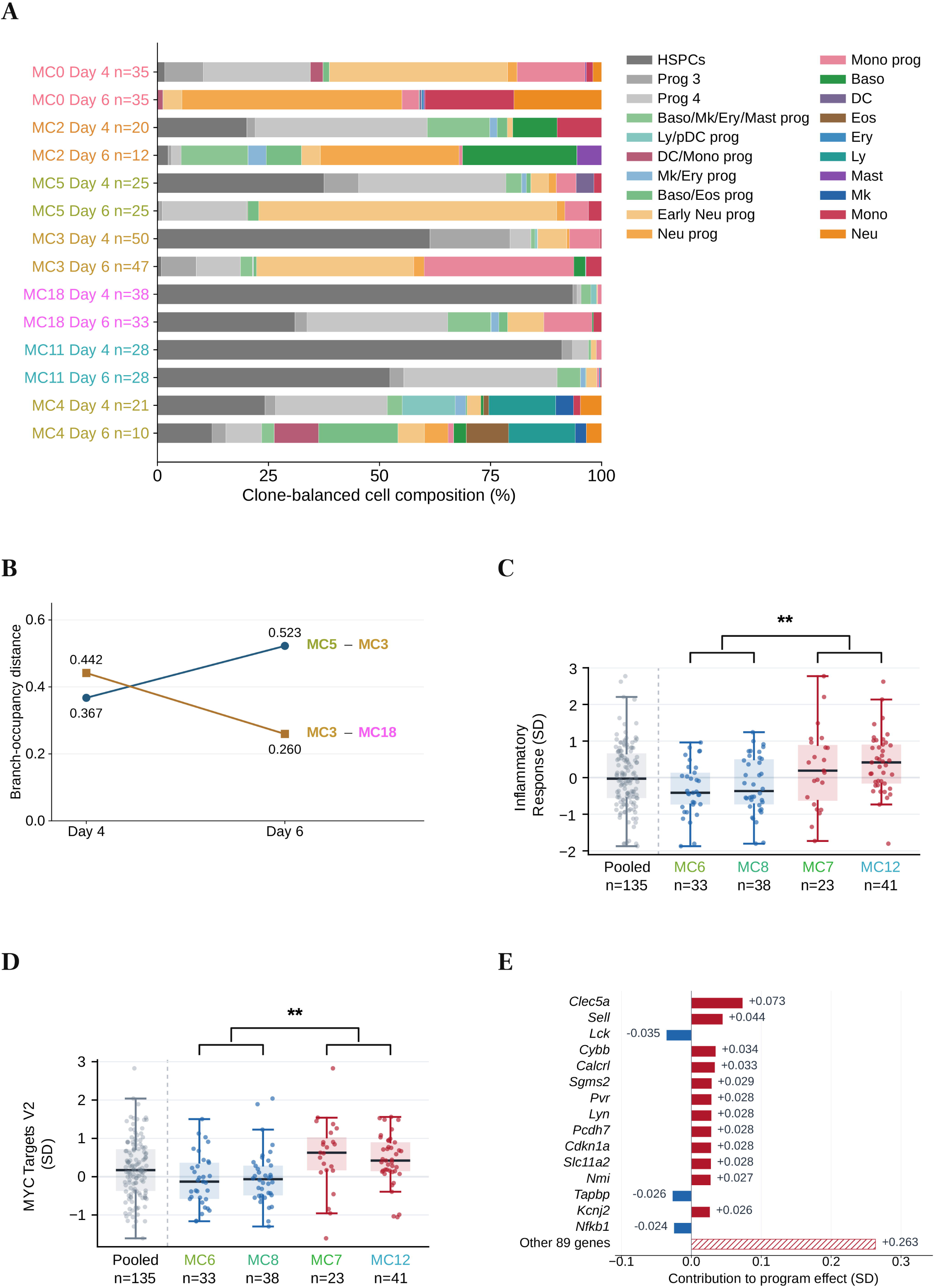
Multilineage metaclones differ in lineage output over time, and early HSPC programs differ between metaclones with distinct clonal dynamics. A. Cell-type proportions at Days 4 and 6, calculated for each clone and averaged over the clones of each metaclone, with HSPCs included. c, number of clones. B. For each metaclone, the fraction of each clone’s cells on the neutrophil, monocyte and basophil branches was averaged over clones, and the plot shows the Euclidean distance between two metaclones on each day (0, identical). MC5 and MC3 became less similar from Day 4 to Day 6, and MC3 and MC18 more similar. Full compositions are in Table S2. C-D. Hallmark scores of Day-2 HSPCs for each clone (adjusted for sequencing library) in the monocyte-output metaclones, shown pooled and by metaclone. Dots are clones; boxes show the median and interquartile range (IQR), and whiskers extend to 1.5 × IQR. Blue, slower-maturing metaclones (MC6, MC8); red, faster-maturing metaclones (MC7, MC12). ** q < 0.05, fast versus slow metaclones, with the two metaclones on each side weighted equally. These two gene sets showed the largest differences; all 50 are shown in Figure S3A and Table S3. C. Hallmark Inflammatory Response. D. MYC Targets V2. E. Contribution of individual genes to the fast-minus-slow difference in the Inflammatory Response score (SD). Red, higher in faster-maturing metaclones; blue, lower. The last bar is the sum of the remaining 89 genes in the set.

Across Palantir^15^-defined trajectories, we compared metaclones with similar lineage output, and then calculated pairwise Euclidean distances between branch occupancy. Profiles showed dynamic divergence (MC5 vs. MC3 distance increased from 0.367 at Day 4 to 0.523 at Day 6) and convergence (MC3 vs. MC18 decreased from 0.442 to 0.260) (Figure 3B). Because the trajectories overlap, occupancies are not mutually exclusive and similar patterns could be seen in the neutrophil and monocyte branches (Table S2).

To investigate whether molecular signatures correlating with metaclone-specific temporal dynamics can be identified in early progenitors, we evaluated Day-2 HSPCs across monocyte producing metaclones. Comparing “fast-maturation” (MC7/MC12) and “slow-maturation” (MC6/MC8) metaclones revealed significant enrichment of Hallmark Inflammatory Response (+0.584 SD, q = 0.0023) and MYC Targets V2 (+0.551 SD, q = 0.0011) in fast-maturing progenitors (Figures 3C, 3D and S3A). These differences were broad: 30 of the 50 Hallmark gene sets were significantly higher in fast-maturing progenitors (q < 0.05), and none were lower, including sets unrelated to blood such as Spermatogenesis (Table S3). The genes contributing most strongly to the inflammatory score were myeloid genes such as Clec5a, Cybb and Lyn (Figure 3E). Consistent with this, Day-2 cells within the fast-maturing metaclones were already positioned further along the monocyte differentiation trajectory (clone median pseudotime 0.14 versus 0.10; Figure 2H). Thus, the elevated inflammatory response score reflects accelerated myeloid maturation rather than active inflammatory signaling. In addition, fast-maturing progenitors exhibited elevated scores for proliferation-associated programs, including MYC targets, E2F Targets (+0.334 SD, q = 0.013) and G2-M Checkpoint (+0.312 SD, q = 0.025).

### WT reference metaclones resolve hidden, clone specific mutation responses

We next evaluated ChronoClone on a longitudinal mutation-tracing dataset (Dnmt3a-R878H and Npm1c induction in LARRY barcoded HSPCs)^16^ (Figures 4A and 4B). In this study, LARRY-barcoded mouse HSPCs were expanded in culture and profiled at Day 7; the cultures were then split into mock-treated WT controls and mutant conditions. Cre induced Dnmt3a-R878H and Flpo induced Npm1c in their respective experiments. Descendants were profiled at Days 14 and 27. Because sister clones are present in all conditions, we first defined 17 reference metaclones, strictly within WT control clones and transferring these labels to mutant sister clones via shared barcodes, present in the mutated conditions. We then compared each mutant metaclone with its WT counterpart (Figure S4). Across 68 metaclone-level comparisons (17 metaclones x 2 mutations x two time points), ChronoClone resolved 12 maturation shifts and 17 cell-type composition changes (Figures 4C, 4D and 4E; Table S4).

**Figure 4:**
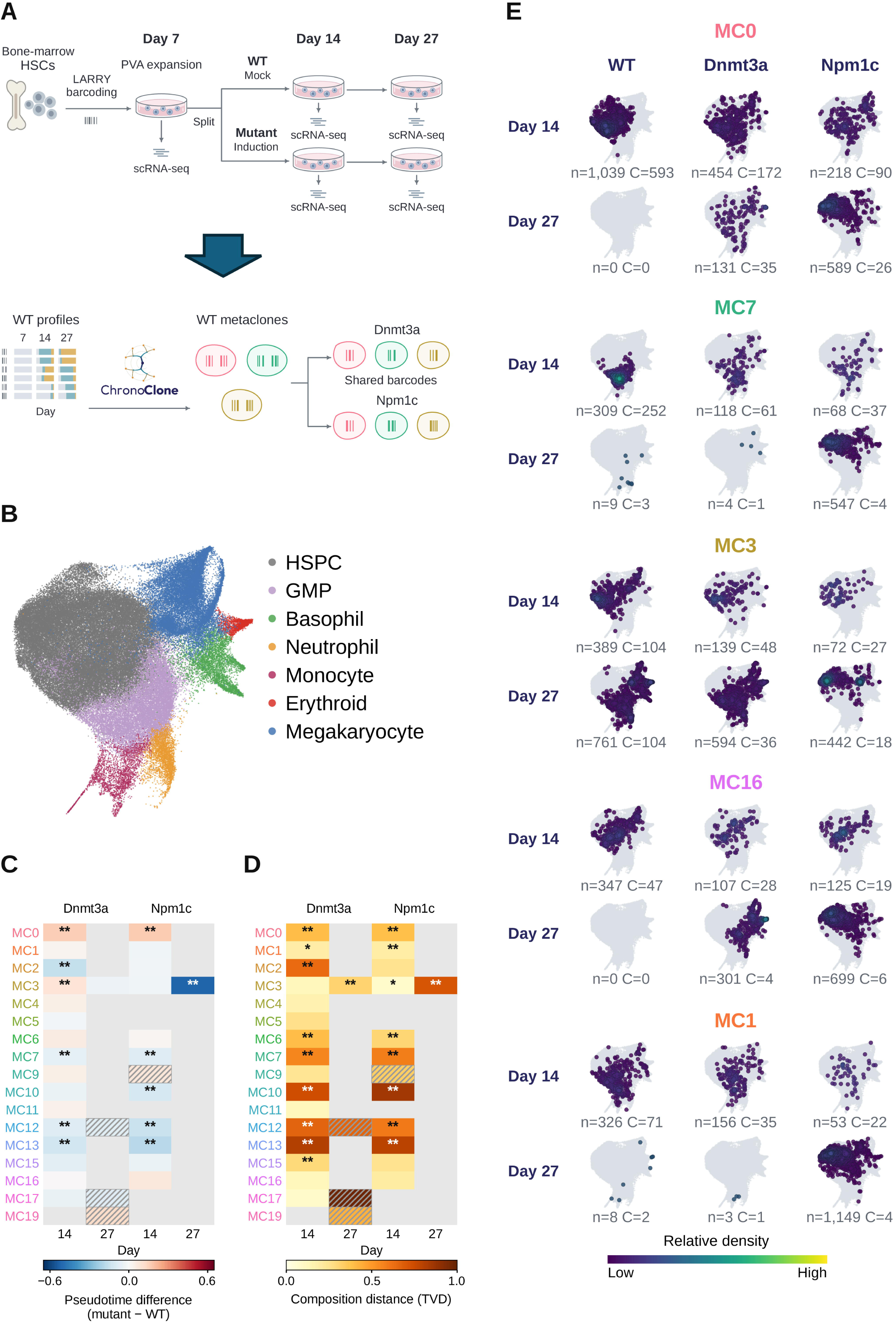
WT metaclones resolve distinct dynamics in a Dnmt3a and Npm1c mutation-tracing experiment. **A.** Bone-marrow HSPCs (Lin^-^Sca1^+^c-kit^+^/Lin^-^ckit^+^) were barcoded with LARRY and expanded in PVA-based HSC expansion media^23^. Day-7 cells were profiled with scRNA-seq, and the cultures were then split into mock-treated WT controls and mutant conditions (Cre for Dnmt3a-R878H and Flpo for Npm1c). Descendants were profiled at Days 14 and 27. Experimental design taken from Singh et al.^16^. **B.** UMAP of all 96,662 cells, with the coordinates and cell-type annotation of the original study (Hematopoietic Stem Cells and Multipotent Progenitors pooled as Hematopoietic Stem and Progenitor Cells - HSPCs)^16^. ChronoClone used the 14,896 cells (1,516 clones) that carry a lineage barcode. **C.** Maturation of mutant clones relative to their WT sister clones in each metaclone: difference in mean pseudotime (mutant minus WT) at Days 14 and 27. ** q < 0.05 (Benjamini–Hochberg correction over 68 tests). Gray, too few cells or clones to estimate; hatched, estimated but too few clones for a test. **D.** As in C, for the change in cell-type composition, measured as the total variation distance between mutant and WT cells (0, identical composition). ** q < 0.05 and * q < 0.10 (a separate Benjamini–Hochberg correction over 68 tests). **E.** Cells of MC0, MC1, MC3, MC7, and MC16 on the landscape in B, for WT, Dnmt3a and Npm1c sister clones at Day 14 (top) and Day 27 (bottom). Color shows relative cell density on a common scale (key); n denotes number of cells and C the number of clones. All metaclones are shown in Figure S4.

Specifically, ChronoClone uncovered three distinct classes of mutant behavior: 1) Lineage skewing, Dnmt3a-induced clones exhibited preferential production of megakaryocytes, erythroid cells, and neutrophils (MC16, MC17, and MC19), a lineage bias absent in both the WT reference baseline and Npm1c-induced clones (Figure S4B). 2) Dynamic latency, Npm1c sister-clones appeared broadly normal at Day 14, but by Day 27 exhibited marked maturation delay, remaining predominantly restricted to the HSPC compartment (MC0, MC1, MC3, MC7, and MC16). 3) Clonal survival recovery, mutant sister-clones persisted at Day 27 despite the complete absence of their corresponding WT counterparts.

## Discussion

Existing computational single-cell lineage inference approaches primarily infer static tree topologies, pairwise transition maps, or continuous cell embeddings. ChronoClone advances this by directly modeling temporal cell-state compositions, grouping clones into data-driven reference metaclones using automated silhouette selection. By shifting focus from terminal endpoint outputs to dynamic trajectory profiles, ChronoClone demonstrates that clones with identical output biases often differentiate at different speeds and produce different lineages at intermediate time points.

Applying ChronoClone to both datasets uncovered dynamic clonal behaviors not reported in the original studies. Weinreb et al.^5^ showed that sister cells differentiate at a consistent pace, they did not evaluate whether differentiation rates vary between distinct clones. Our analysis revealed substantial inter-clonal heterogeneity in maturation speed within the same lineage, showing that the fastest-maturing clones reduced in size by Day 6 while slower-maturing clones continued to expand. Singh et al.^16^ compared mutant and WT sister clones at Day 27, ChronoClone detected mutation driven shifts in maturation as early as Day 14, just one week after induction, skewing differentiation in specific metaclones (MC17, MC19) while delaying it in others (MC0, MC1, MC3, MC7).

ChronoClone can also be used to study perturbations. Conventional comparisons between mutant and WT cells obscure heterogeneous behaviors by pooling clones with divergent behaviors. By defining metaclones within WT cells and then assigning those labels onto matched mutant sister clones, ChronoClone enables controlled comparisons of the effect of the mutation on specific time-resolved behaviors. Applying this strategy to the mutation-tracing dataset revealed context dependent phenotypes. More broadly, the sister-clone mapping approach can be readily applied to study diverse genetic or drug perturbations, provided that barcoded sister clones are sampled and present across both control and perturbed conditions.

ChronoClone does not use gene expression to group clones; it needs only clone identities, sampling times and a state label for each cell. The user chooses what the states are: here they were cell types from scRNA-seq, but they could be cell states scored by live imaging, the expression of a fluorescent reporter, or accessibility of selected peaks in scATAC-seq. ChronoClone can therefore be applied to any longitudinal lineage-tracing experiment, and to tissues other than blood, such as organoids modeling normal and malignant organs. Furthermore, by establishing prospective metaclone groupings, researchers can retrospectively query upstream single-cell transcriptomic or epigenomic datasets to identify early molecular markers predictive of long-term clonal fate.

### Limitations of the study

Several limitations are to be considered when applying of ChronoClone. Firstly, although ChronoClone can be run on two post-initiation time points without sampling the starting population (provided the initial cell type of each clone is known), the resulting metaclones capture dynamics solely across these two time points, and preclude comparisons of early progenitor states across divergent trajectories. ChronoClone has only been benchmarked on datasets comprising of an initial progenitor time point followed by three subsequent time points, therefore sampling across three time points is strongly recommended. Second, reliable estimation of state composition requires sufficient cell numbers per clone at each time point, and the overall temporal resolution is limited by the frequency and spacing of the experimental sampling. Third, because clones are represented as time-resolved proportions of annotated cell states, metaclone resolution depends directly on the accuracy and granularity of cell-type annotations. Overly coarse annotations may obscure subtle dynamic differences, whereas overly fine-grained annotations may split a single behavior across multiple metaclones. Finally, novel perturbation induced cell states absent in the WT baseline cannot be captured within the reference metaclone behaviors.

A separate limitation concerns clonal recovery. The absence of a clone at a given time point could be due to technical barcode dropout or insufficient numbers of cells being sequenced. Finally, although the silhouette score automates partition of metaclones, the resulting metaclones are still influenced by the chosen range of clustering resolution and tree-cut criteria.

## Supporting information

Supplemental Table Guide

## Resource availability

### Lead contact

Lead contact Melania Barile and Nicola Wilson.

### Materials availability

This study did not generate new unique reagents.

### Data and code availability

Both datasets analyzed in this study are published and publicly available. The lineage-tracing data of Weinreb et al. are in GEO under accession GSE140802 (in vitro sample GSM4185642), and the cell-state landscape and annotations used here are taken from Gao et al.^10^ The mutation-tracing data of Singh et al. are in GEO under accession GSE266232, with the original analysis code and objects at https://doi.org/10.5281/zenodo.14261507. Accession numbers and links are also listed in the key resources table. This study did not generate new data.

The ChronoClone package is available at https://github.com/Gottgens-lab/ChronoClone. All code used to analyze the data and generate the figures, including the executed notebooks and the tables behind each panel, is available at https://github.com/Gottgens-lab/ChronoClone_paper. Both repositories will be public as of the date of publication, and the released versions will be archived with DOIs listed in the key resources table.

Any additional information required to reanalyze the data reported in this paper is available from the lead contact upon request.

## Acknowledgments

Work in the Göttgens laboratory is supported by Wellcome (206328/Z/17/Z and 309075/Z/24/Z), Medical Research Council (MR/W031663/1) and Aging Biology Foundation. Hiromitsu Harimoto’s doctoral studies were supported in part by a scholarship from Rotary International. Weizhong Zheng was supported by BBSRC Impact Acceleration Account. This research was funded in whole, or in part, by the Wellcome Trust [203151/Z/16/Z, 203151/A/16/Z and 226795/Z/22/Z] and the UKRI Medical Research Council [MC_PC_17230]. For the purpose of open access, the author has applied a CC BY public copyright license to any Author Accepted Manuscript version arising from this submission.

## Author contributions

H.H performed all data analysis and wrote the paper. W.Z provided criterial knowledge for simulation testing. M.B supervised and performed computation analysis. All authors critically reviewed and approved the final manuscript. B.G. M.B and N.K.W conceived, conceptualized and supervised the study.

## STAR Methods

### Key resources table

| Reagent or resource | Source | Identifier |
|---|---|---|
| Mouse hematopoietic lineage-tracing data | Weinreb et al.<sup>5</sup> | GEO: GSE140802; in vitro sample GSM4185642 |
| Hematopoietic state landscape and annotations | Gao et al.<sup>10</sup> | CLADES; https://doi.org/10.6084/m9.figshare.27908142.v2 |
| Processed Weinreb analysis object and figure-source tables | This paper | ChronoClone_paper repository; versioned release pending |
| Ex vivo mutation-tracing data and source analysis code | Singh et al.<sup>16</sup> | GEO: GSE266232; https://doi.org/10.5281/zenodo.14261507 |
| ChronoClone and manuscript analysis notebooks | This paper | https://github.com/Gottgens-lab/ChronoClone; https://github.com/Gottgens-lab/ChronoClone_paper |
| Hallmark gene sets | Liberzon et al.<sup>17</sup> | Fixed MSigDB Hallmark 2020 resource |
| Palantir | Setty et al.<sup>15</sup> | https://github.com/dpeerlab/Palantir |
| Scanpy | Wolf et al.<sup>18</sup> | https://github.com/scverse/scanpy |
| scikit-learn | Pedregosa et al.<sup>19</sup> | https://scikit-learn.org |
| SciPy | Virtanen et al.<sup>20</sup> | https://scipy.org |
| statsmodels | Seabold and Perktold<sup>21</sup> | https://www.statsmodels.org |
| UMAP | Becht et al.<sup>22</sup> | https://github.com/lmcinnes/umap |
| Python analysis environments and package versions | This paper | Exact versions and execution commands in the executed notebooks |

### Experimental model and study participant details

No new mouse or human material was generated; analysis was performed on published datasets. Weinreb et al.^5^ barcoded mouse hematopoietic progenitors with LARRY and cultured them in vitro, sampling at Days 2, 4 and 6. We used the cell-state landscape and annotations described in CLADES^10^. Singh et al.^16^ barcoded mouse HSPCs, expanded them in PVA-based cultures^23^, profiled Day-7 cells, then split the cultures into mock-treated WT controls and mutant conditions (Cre for Dnmt3a-R878H and Flpo for Npm1c). Descendants were profiled at Days 14 and 27. We used the landscape, cell-type annotation and pseudotime as defined in the original publication, with Hematopoietic Stem Cells (HSCs) and Multipotent Progenitors (MPPs) pooled as hematopoietic stem and progenitor cells (HSPCs) throughout this paper. Mice, culture, barcoding and sequencing are described in the original studies.

### Method details

ChronoClone takes as input a clone label, a cell-state label and a time point for every cell, together with one or more starting cell types that contain the originally labeled cells, the most primitive progenitors of the clones. A clone is retained when all of its cells at the first time point belong to these starting cell types. For each clone and time point, cells are counted per state and converted to proportions.

Clones are clustered on a nearest-neighbor graph of their profiles with the Leiden algorithm over a range of resolutions. The fraction of resolutions in which two clones fall in the same cluster defines a consensus distance between them; hierarchical clustering of this distance is cut at the threshold giving the highest mean silhouette coefficient across all clones, and the resulting groups are called metaclones. The number of metaclones is therefore not set in advance.

When an experiment has several conditions, metaclones are defined in a reference condition only (here, the WT control). A clone in another condition takes the metaclone label of the WT clone with the same LARRY barcode; the clustering is not rerun on the other conditions. Parameter values and seeds are detailed in the GitHub repository (https://github.com/Gottgens-lab/ChronoClone_paper).

### Clone-level statistics

Every cell-level measurement is first averaged within its clone, and every clone then contributes equally to its metaclone, so that a few large clones cannot dominate a metaclone. Conditions are compared within a sequencing sample, and the per-sample differences are averaged across samples. Each sample is weighted by the smaller of its WT and mutant clone counts, so that a sample in which one condition is barely represented contributes little. Two quantities are compared: maturation, i.e. the mutant-minus-WT difference in mean pseudotime, and composition, the total variation distance between the mutant and WT cell-type proportions. A comparison is made only when there are enough WT and mutant cells and clones.

Cells of the same clone are not independent, and WT and mutant sister clones share an origin, so clones rather than cells are treated as independent units: standard errors are clustered by clone, and P values come from a multiplier bootstrap with one random weight per clone^24^. Maturation and composition are corrected separately by the Benjamini–Hochberg procedure^25^ across the 68 planned comparisons (17 metaclones × 2 mutations × 2 days), and the branch analysis across all 340 of them (those comparisons × 5 branches).

### Simulation benchmark

To test how well ChronoClone recovers known groups of clones, we simulated clonal data from a compartment model in which each of 12 time-resolved programs sets the proliferation, differentiation and loss rates of 13 cell states. The programs generated 517 clones, each sampled at four time points. We then distorted the last three time points in two ways: Gaussian count variation, and random state redistribution (random reassignment of cells between states). Both noise levels varied from 0% to 100%, with 100 repeats per combination. ChronoClone, CLADES preprocessing, K-means and Ward clustering were tested on the data, and accuracy was assessed as the Adjusted Rand Index (ARI) ^26^ against the simulated programs. clone2vec was not benchmarked because it requires expression neighborhoods, which this count-based generator does not produce.

### Murine adult hematopoiesis dataset: metaclones, lineage output and branches

Clone profiles used Days 2, 4 and 6 with Day-2 HSPCs as the starting cell state, giving 20 metaclones over 569 clones and 7,733 cells. Differentiation progression was assessed with pseudotime coordinates as calculated via Palantir^15^, run from a Day-2 HSPC root with megakaryocyte, basophil, neutrophil, mast-cell and monocyte terminal states. Cells were assigned to branches by Palantir according to their probability of reaching each terminal state, so a cell could be assigned to more than one branch, as early cells can still reach several terminal states^15^. The lineage output of a metaclone is the composition of its differentiated cells pooled over the three days, and branch occupancy is the fraction of a clone’s cells lying in a branch, averaged over the total number of clones within the metaclone. To test whether metaclones with the same lineage output progressed along a branch at different rates, each clone was summarized by the median pseudotime of its cells on that branch, and clone medians were compared between metaclones with a Kruskal–Wallis test on each day; metaclones with fewer than three clones on the branch that day were excluded. The 12 tests (neutrophil- and monocyte-output metaclones on their own branch at Days 2, 4 and 6, and multilineage metaclones on the neutrophil, monocyte and basophil branches at Days 4 and 6) were corrected together with the Benjamini–Hochberg procedure (Table S2). The number of sequenced cells per clone was compared between the fast-maturing (MC7, MC12) and slow-maturing (MC6, MC8) monocyte-output metaclones on each day with a two-sided Mann– Whitney test, with clones as the units. Because all clones were sampled in the same way, these counts reflect relative rather than absolute clone sizes.

### Day-2 HSPCs clone-level transcriptomic analysis

scRNA-seq profiles of individual Day-2 HSPCs were linked to their clones via shared cell barcodes. Because the deposited expression is log-normalized, counts were recovered per cell, summed within each clone, and scored for the 50 Hallmark programs, giving one score per clone and program in standard deviations^17^. Metaclones were compared by ordinary least squares with each clone’s sequencing-library composition as a covariate and clone-level heteroskedasticity-consistent (HC3) standard errors^27^, correcting one-versus-rest and pairwise comparisons in separate Benjamini–Hochberg families. The comparison of the “fast-maturation” monocyte metaclones (MC7, MC12) with the “slow-maturation” ones (MC6, MC8) gives the two metaclones on each side equal weight and is corrected as its own family of 50 tests. Each effect was recalculated after removing each sequencing library in turn, after accounting for HSPC state composition, and with size factors that exclude mitochondrial genes; all versions are in Table S3.

### Comparing mutated sister-metaclones with WT metaclone reference

Each mutant clone retained the metaclone label of its WT sister-clone, mutant and WT descendants were then compared within every metaclone, mutation and day, using the summaries and statistics described above. The unit of each comparison is a clone within one sequencing sample, carrying its mean pseudotime and proportions across seven cell types (HSPCs, GMPs, and megakaryocyte, erythroid, basophil, monocyte and neutrophil progenitors). Branch comparisons repeat the same procedure using only the cells on each of the five Palantir branches (megakaryocyte, erythroid, basophil, monocyte and neutrophil), with each clone’s mean pseudotime and the numbers of cells and clones recalculated within that branch. Because a cell can lie on more than one branch, and each mutation is compared with WT separately, these are not tests between branches or between mutations. The direction of the Day-27 Npm1c difference in MC3 was confirmed by resampling whole clones. Figure S4 shows where the cells from each metaclone lie on the transcriptomic landscape. Descriptive cell-type proportions in Table S5 were averaged first over clones within each sample and then over samples. Table S4 reports the weighted mutant–WT comparisons.

## Supplemental information

**Figure S1:**
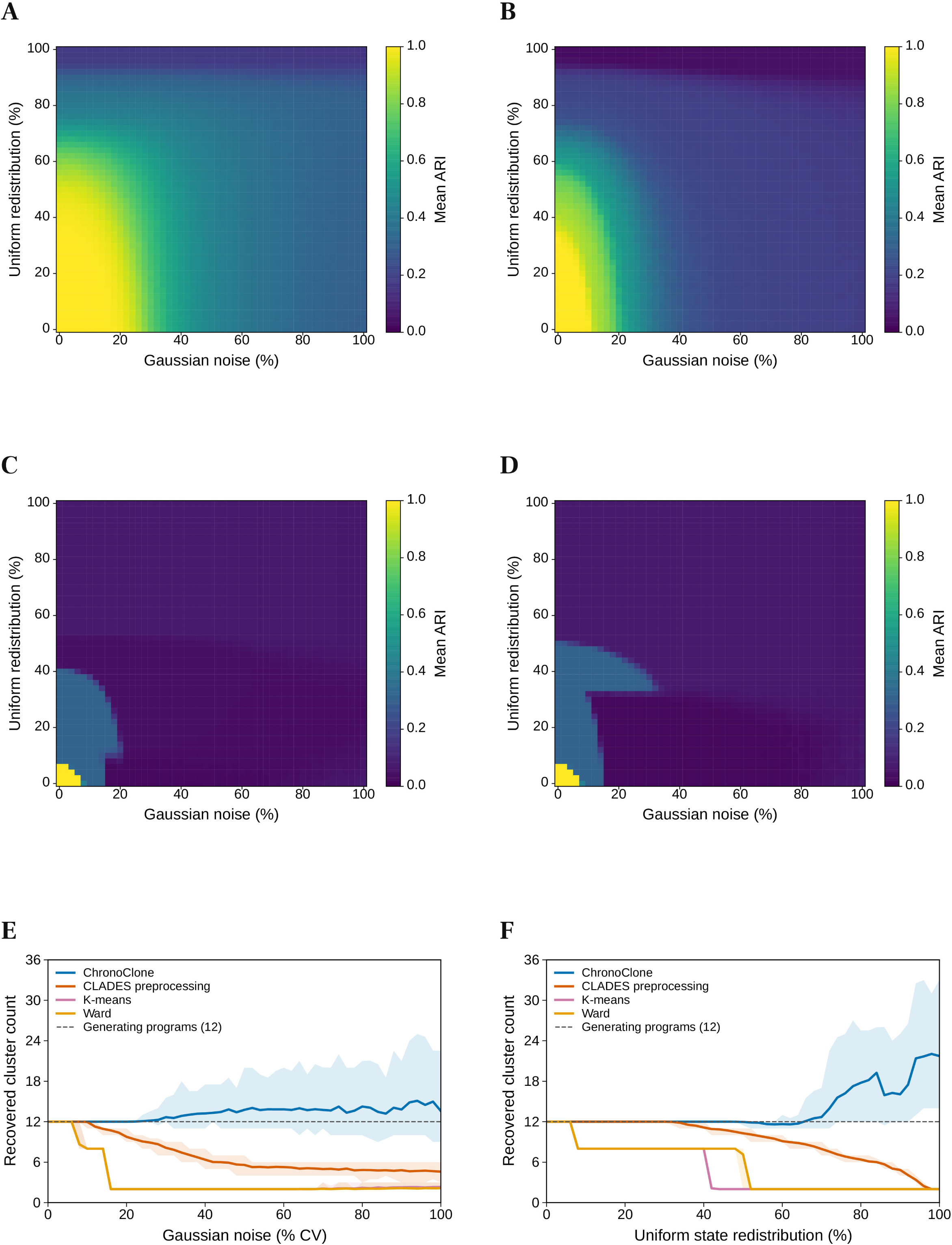
ChronoClone demonstrates high noise robustness in ODE simulations. **A.** ChronoClone accuracy (mean ARI over 100 repeats) when Gaussian noise and random state redistribution are combined at 51 levels each, related to Figure 1. Noise is applied at t2–t4; t1 is unchanged. A–D use the same 0–1 color scale. **B.** As in A, for CLADES preprocessing. **C.** As in A, for K-means, with the number of clusters chosen by silhouette score. **D.** As in A, for Ward clustering, with the number of clusters chosen by silhouette score. **E.** Number of groups found by each method as Gaussian noise increases, without state redistribution. Lines show the mean and ribbons the 2.5th–97.5th percentiles of 100 repeats. The dashed line marks the 12 simulated programs. **F.** As in E, for random state redistribution without Gaussian noise.

**Figure S2:**
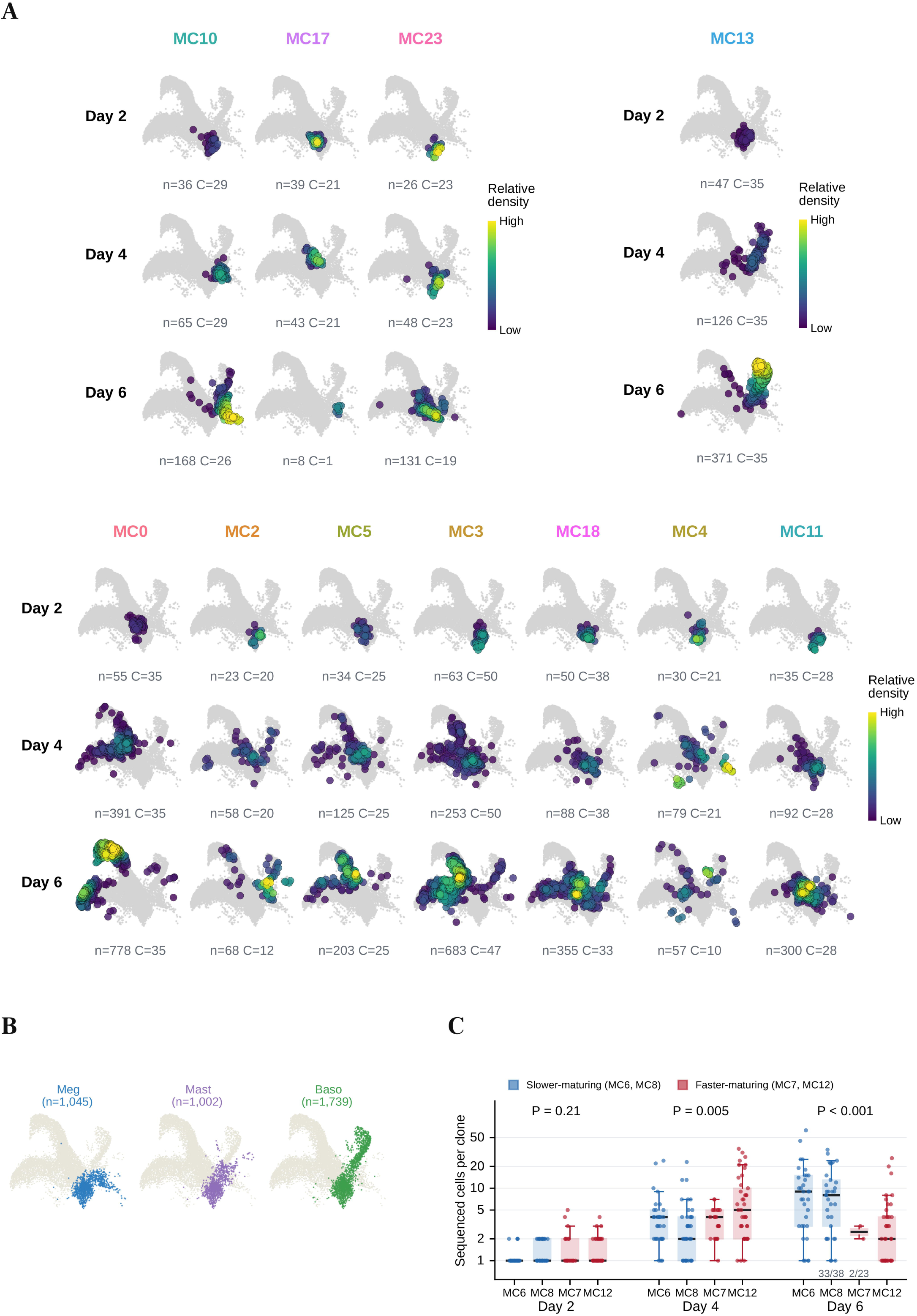
ChronoClone uncovers dynamic differences within metaclones with similar lineage outputs. **A.** The remaining metaclones of the Weinreb et al. dataset^5^, shown as in Figure 2D. Together with Figure 2D, all 20 metaclones are shown. **B.** Megakaryocyte, mast-cell and basophil trajectories, shown as in Figure 2C; c denotes number of cells. **C.** Sequenced cells per clone on each day for the slower-maturing (blue; MC6, MC8) and faster-maturing (red; MC7, MC12) monocyte-output metaclones. Only clones recovered on that day are shown; numbers below a box give recovered/total clones where some were not recovered. Dots are clones; boxes show the median and interquartile range (IQR), and whiskers extend to 1.5 × IQR. P, two-sided Mann–Whitney test of fast versus slow clones, counting unrecovered clones as zero. Because all clones were sampled in the same way, the counts reflect relative clone size.

**Figure S3:**
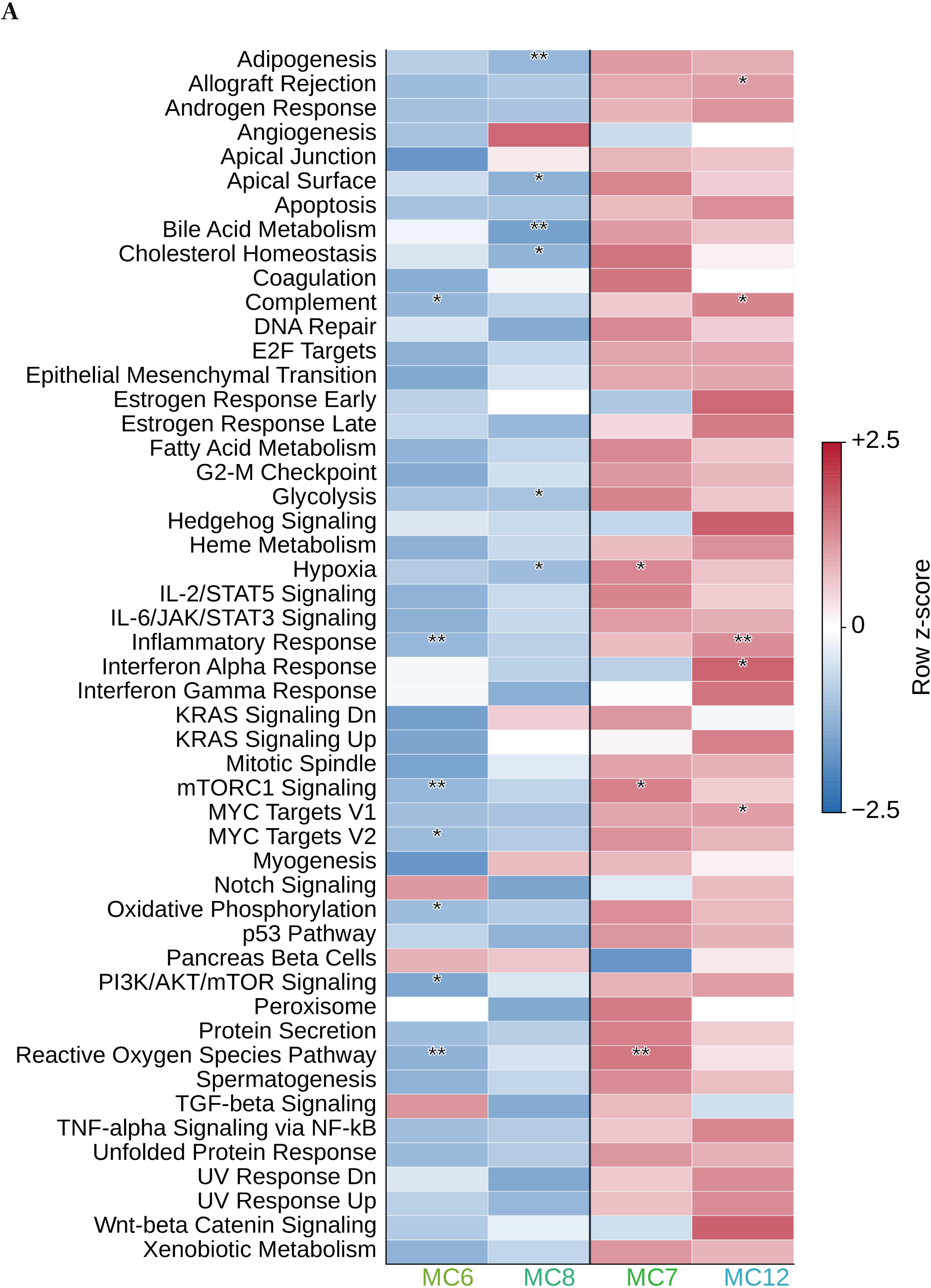
Early HSPC programs differ between metaclones with distinct clonal dynamics. **A.** All 50 Hallmark gene sets for the monocyte metaclones, related to Figure 3. Color shows the row z-score of the library-adjusted metaclone means. ** q < 0.05 and * 0.05 ≤ q < 0.10, one metaclone versus the rest (Benjamini–Hochberg correction over 950 tests).

**Figure S4:**
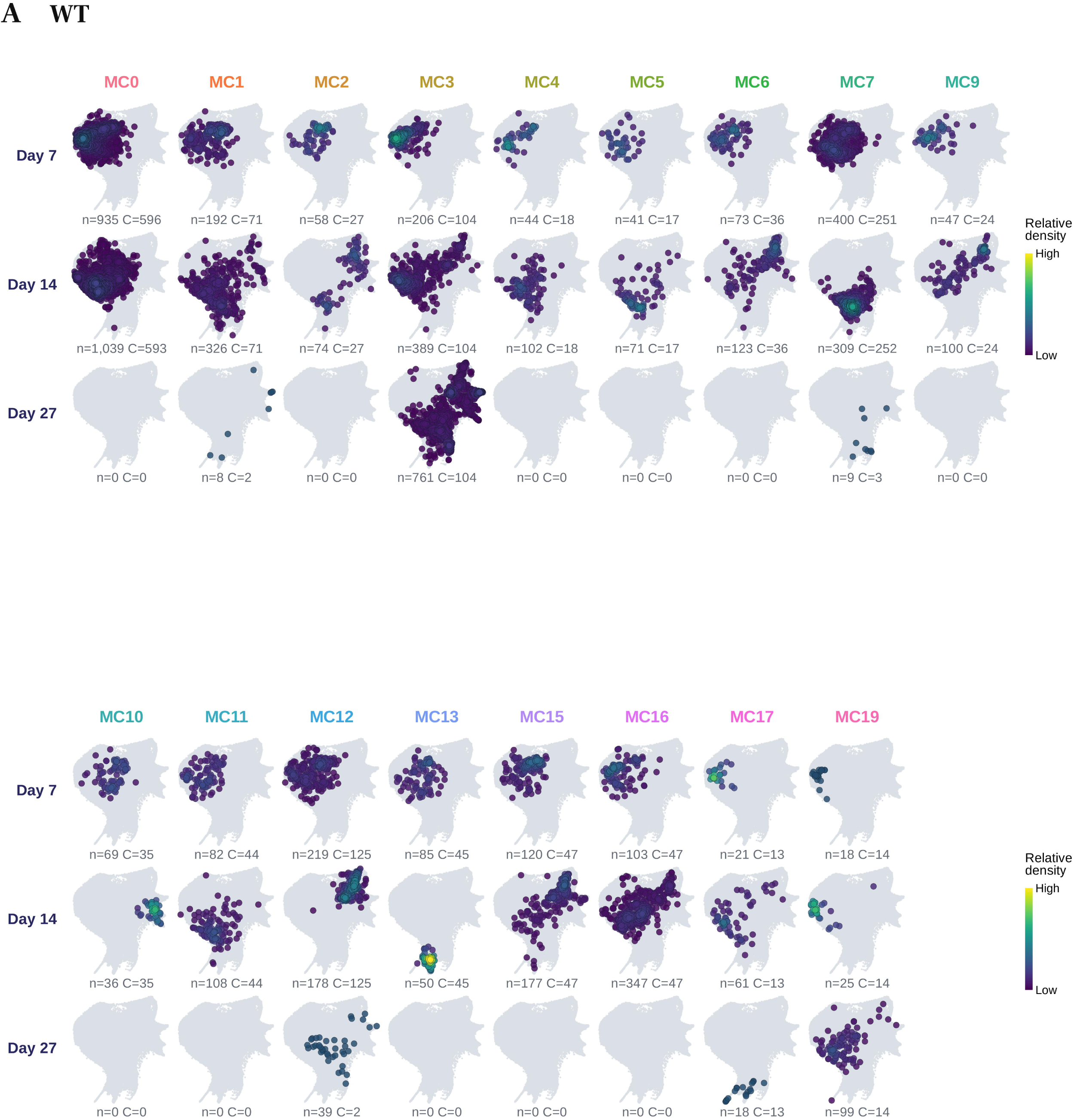
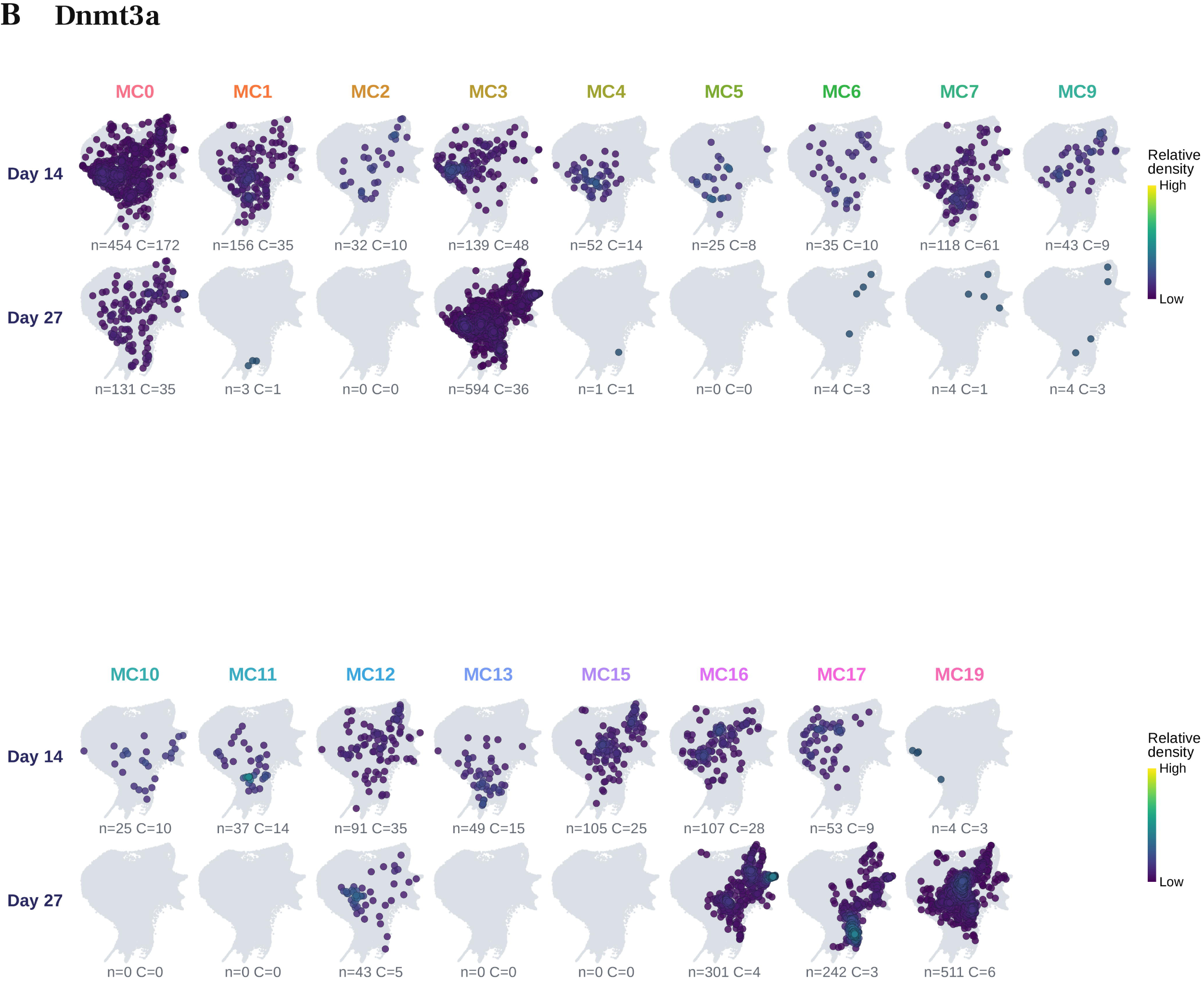
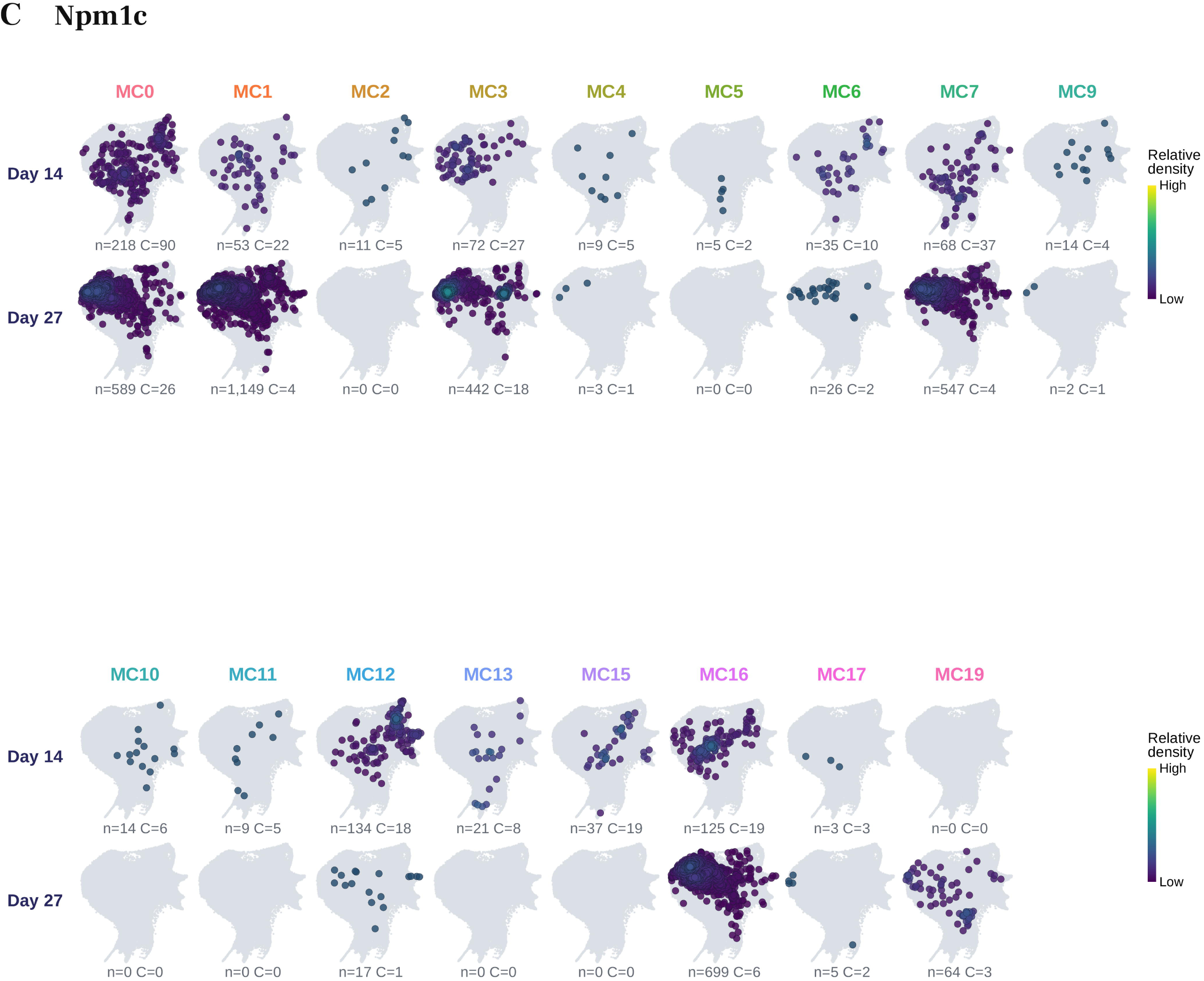
WT metaclones resolve distinct dynamics in a Dnmt3a and Npm1c mutation-tracing experiment. **A.** Distribution of WT cells on the transcriptomic landscape for each of the 17 reference metaclones (columns) at Days 7, 14 and 27 (rows). Gray, no cells recovered. **B**. As in A, for the Dnmt3a sister clones. The Day-27 lineage bias of MC16, MC17 and MC19 is descriptive, because too few WT clones remained for a formal test. **C**. As in A, for the Npm1c sister clones. Day 7 (before induction) is shown only in A. In the density maps, each point is a cell on the landscape in Figure 4B; density is scaled within each metaclone, condition and day, with the same settings for all maps. Maps with fewer than 20 cells or three clones show individual cells (navy points). n denotes number of cells and C the number of clones.

Table S1. ODE simulation design and recovery of the simulated programs.

Table S2. Clonal dynamics, lineage trajectories and barcode linkage in the Weinreb dataset.

Table S3. Gene programs in Day-2 HSPCs from Weinreb metaclones.

Table S4. Mutant–WT comparisons within Singh reference metaclones.

Table S5. Cell-state landscapes, WT metaclone profiles and clone recovery in the Singh dataset.

Table S6. NMF programs in Day-7 HSPCs from the Singh dataset.

