## Supplemental Table Guide for "ChronoClone links temporal clonal behavior to lineage output and mutation responses"

ChronoClone supplementary tables

The six archives contain data for the simulation benchmark and the lineage-tracing studies by Weinreb et al. and Singh et al. Unzip the relevant archive and open contents.csv to see what each file contains and what one row represents. Use columns.csv for field definitions. The data files are in the data folder. Import clone and cell identifiers as text.

### ODE simulation design and benchmark recovery

**Table_S1.zip** Compare the four methods across noise levels in primary_recovery.csv, which reports the adjusted Rand index and the number of groups recovered. single_noise_slices.csv gives the results when one noise parameter varies at a time.

### Weinreb metaclone dynamics and barcode linkage

**Table_S2.zip** Use Multilineage_composition.csv to compare cell-type proportions across the seven multilineage metaclones. Pseudotime_tests.csv gives the within-day tests of differences between metaclones along each lineage branch.

### Day-2 HSPC gene programs in Weinreb metaclones

**Table_S3.zip** Find the inflammatory response and MYC comparisons from Figures 3C and 3D in S3_Selected_Hallmark_effects.csv. S3_Hallmark_effects.csv gives the full set of Hallmark comparisons. Additional files report clone-level program usage from consensus nonnegative matrix factorization (cNMF).

### Mutant–WT comparisons within Singh reference metaclones

**Table_S4.zip** Use state_responses.csv for changes in maturation and composition_responses.csv for changes in cell-type proportions. Both report comparisons of mutant descendants with their WT sisters within each metaclone and day.

### Singh cell-state landscapes, WT metaclone profiles and sampled recovery

**Table_S5.zip** Find the WT profiles at Days 7, 14 and 27 in wt_reference_profiles.csv. Check how many cells and clones were observed in each condition and day using all_snapshot_support.csv.

### NMF programs in Singh Day-7 HSPCs

**Table_S6.zip** Use S6_Program_annotations.csv to identify the genes supporting each of the eight programs. S6_MC_usage_intervals.csv reports mean relative program usage by metaclone and intervals obtained by resampling clones.
